# Reliability of Spatiotemporal Neuromuscular Activity Patterns in Magnetomyography Across Force Levels

**DOI:** 10.64898/2026.05.24.727496

**Authors:** Haodi Yang, Burak Senay, Benedict Kleiser, Justus Marquetand

**Author notes:** Corresponding Author: Prof. Dr. Justus Marquetand Pfaffenwaldring 5a, 70569 Stuttgart, Germany Phone: +497071/2981196, Fax +497071/294488.

## Abstract

**Background:** Magnetomyography (MMG) using an optically pumped magnetometer (OPM) provides a contactless and non-invasive approach to assess neuromuscular activity. However, given the limited studies using optically pumped magnetometer magnetomyography (OPM-MMG) and that those primarily used a single OPM, it remains to be characterized how stable individual neuromuscular activity patterns are over time and across different force levels when using multiple OPMs, specifically an array-based MMG.

**Methods:** 12 healthy subjects performed ramped isometric contractions at 10, 20, 40, 60, and 80% maximal voluntary contraction (MVC), while a 2×3 OPM array recorded MMG signals from the tibialis anterior (TA). For the test-retest reliability, ramps at 20% and 60% MVC were each repeated once. For each subject, Spearman’s rank correlation was computed across all OPM sensors between force conditions to assess consistency of feature rankings, and within-subject spatial repeatability between repeated 20% and 60% MVC ramps was quantified using ICC (3,1) (two-way mixed-effects, single-measure, absolute agreement).

**Results:** In 9 of 12 subjects, Spearman’s rank analyses showed generally high correlations across force levels (ρ = 0.37 to 0.91, p<0.05), whereas in the remaining 3 subjects, spatial patterns were less stable (ρ = −0.73 to 0.58, p<0.05). ICC (3,1) between repeated ramps indicated high within-subject spatial repeatability of the array pattern in 9 out of 12 subjects, with ICC > 0.75 at 20% MVC (5 out of 9 subjects) and 60% MVC (6 out of 9 subjects), respectively. In contrast, the remaining 3 subjects showed lower ICCs (20% MVC: −0.28 to 0.51; 60% MVC: 0.22 to 0.80).

**Conclusion:** Array-based OPM-MMG shows high within-subject stability of spatial patterns across force levels and strong test–retest repeatability in the majority of subjects (9 out of 12), supporting its use for characterizing force-dependent neuromuscular activity while acknowledging inter-subject variability.

## 1. Introduction

Biomagnetism research has advanced significantly with the advent of the superconducting quantum interference device (SQUID) [1,2], which has been widely applied over the past few decades to record human magnetic signals, such as magnetoencephalography (MEG) [3–5] and magnetocardiography (MCG) [6,7]. However, the field of application towards skeletal muscles, i.e., magnetomyography (MMG) [8–11], has been relatively limited. This is primarily because skeletal muscles are widely distributed throughout the body and exhibit considerable variation in size, depth, and anatomical structure, which consequently poses complexities for optimal sensor placement for MMG, as SQUID is often inflexible and bulky due to their need for cryogenic cooling (i.e., −269 ◦C for liquid helium) [12–15]; these practical factors greatly limited the use of SQUID for MMG, thus hindering its research.

With recent technological advancements, the optically pumped magnetometer (OPM, quantum sensors based on the Zeeman effect) has emerged as a high-sensitivity (10−20 ft [3,13,16]), a non-invasive and contactless option for measuring MMG signals [17]. Compared to the conventional SQUID system, OPM operates at room temperature, eliminating the need for complex cooling systems, and the smaller size allows for more flexible placement near the target muscle and can even be used within portable magnetic shields [18]. Consequently, OPMs have made it highly feasible to conduct MMG research. While surface electromyography (sEMG) [19] — the non-invasive gold standard for recording neuromuscular electrical activity — requires skin preparation and electrode attachment, which can compromise signal quality due to skin impedance, contact issues, and motion artifacts, optically pumped magnetometer-based MMG (OPM-MMG) provides a completely contactless and non-invasive alternative that addresses these limitations. Therefore, OPM-MMG is emerging as a promising contactless approach for assessing neuromuscular function.

Although OPM-MMG has demonstrated its potential in neuromuscular research, most recent studies have used only a single commercial OPM sensor for MMG signal recording [8,9,20]. While a single sensor is easy to position, its coverage of the muscle area is limited, reflecting only the localized magnetic neuromuscular information of the target muscle and not adequately characterizing the spatial activity patterns of the entire muscle across different recording locations. This limitation is particularly relevant, as voluntary force is generated by the recruitment and firing rates of motor units — the smallest functional units under neural control — whose fibers are distributed across distinct territories within the muscle volume and are often non-uniform [21]. Consequently, changes in motor unit recruitment generate region-specific activation patterns that cannot be adequately resolved from a single recording position. As evidenced by sEMG studies with multiple electrodes in an array configuration, i.e., high-density surface electromyography (HDsEMG) studies, such spatial information can be used for motor unit decomposition, biofeedback, and prosthetic control [22–24]. Therefore, going beyond single OPM-MMG towards an array-based OPM-MMG approach offers not only broader spatial coverage but also the potential to characterize the physiological organization of muscle activation. Despite this potential, it remains unclear whether spatial MMG activity patterns are stable within subjects, repeatable across consecutive measurements, and consistently preserved or reorganized across varying target force levels.

To investigate these unresolved questions, the present study systematically evaluates the stability of spatial MMG activity of the tibialis anterior (TA) muscle in 12 healthy subjects during isometric contractions at different target force levels (10%, 20%, 40%, 60%, and 80% of maximal voluntary contraction, MVC) using a 2 × 3 OPM sensor array. In addition, the repeatability of the tasks was examined by conducting the 20% and 60% MVC trials twice. The study focuses on two scientific questions: first, whether the spatial patterns of MMG activity recorded by the OPM array exhibit within-subject stability and repeatability; second, whether consistency and variability in spatial patterns exist across different subjects. We hypothesize that the majority of subjects will display relatively stable spatial activity patterns across varying force levels and repeated tests, while a subset of subjects may exhibit higher spatial variability, which could be associated with individual-specific motor unit recruitment strategies.

## 2. Materials and methods

### 2.1 Subjects

Twelve subjects volunteered for the study (five females and seven males; mean age: 29.25 ± 3.91 years; mean body mass index: 22.80 ± 2.51 kg/m^2^). All subjects gave informed consent before participating in the study. The study was conducted in accordance with the Declaration of Helsinki and approved by the ethics committee of the University of Tübingen.

### 2.2 Muscle Ultrasound and OPM Array

Before the recordings, a board-certified sonographer (JM) used an ultrasound system (Mindray TE7, 14-MHz linear probe) to accurately localize the left TA to be voluntarily activated during the study’s paradigm (i.e., the left TA) and to quantify fat thickness. Based on ultrasound results, the TA area was marked on each subject’s skin. Given intersubject variability in TA anatomy, morphology, and size, an anatomical normalization procedure was applied to define a comparable recording region across subjects: specifically, transverse tangent lines (perpendicular to the TA’s longitudinal axis) were drawn at the proximal and distal ends of the TA. Based on these two tangents, the TA muscle was equally divided into three segments: proximal, middle, and distal (Figure 1A), with the middle segment designated as the target measurement zone.

**Figure 1.**
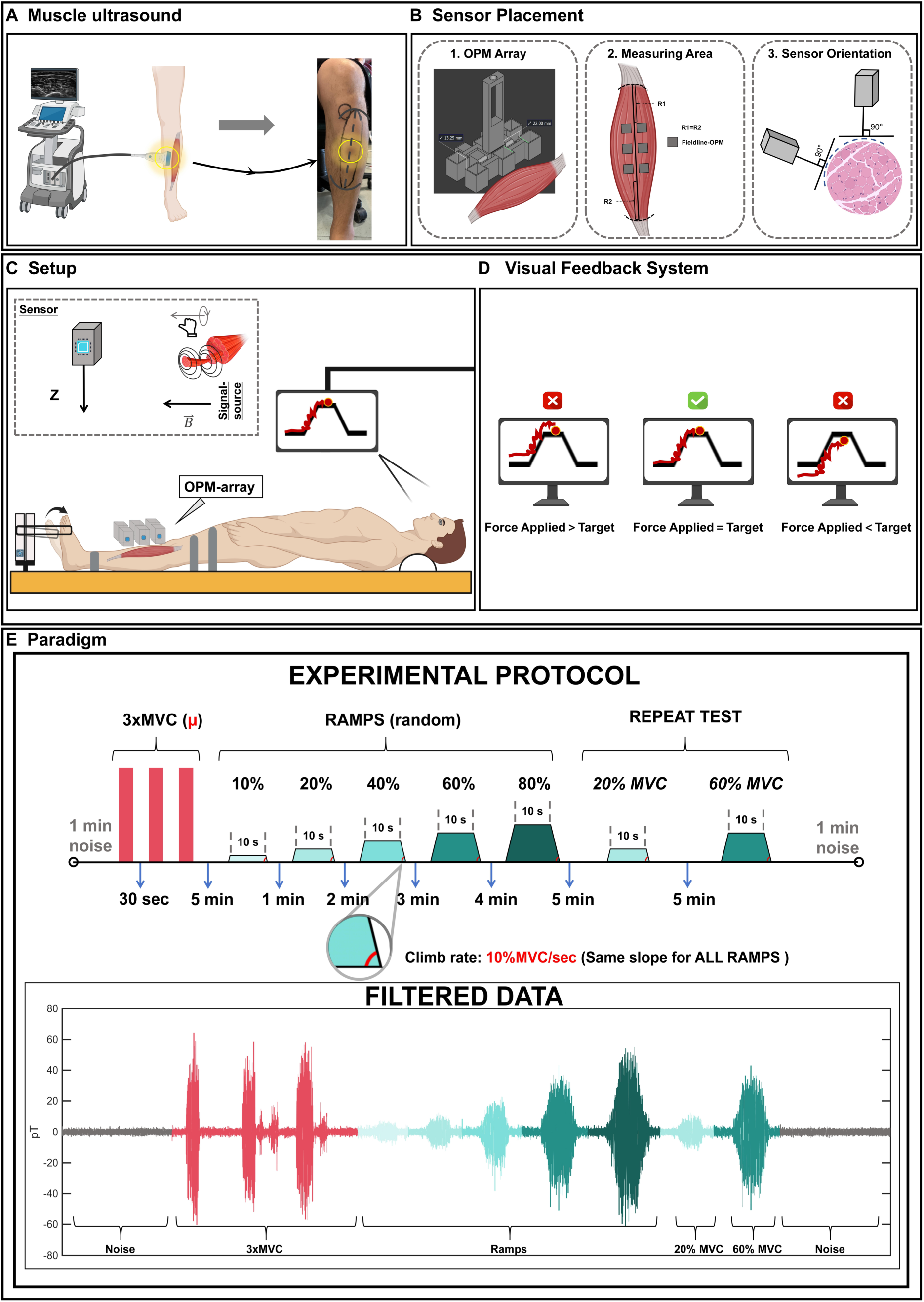
Experimental setup, sensor placement, and measurement protocol for optically pumped magnetometer magnetomyography (OPM-MMG). **(A)** Ultrasound-guided identification of the tibialis anterior (TA) and skin marking of the target recording area. **(B)** Sensor placement and orientation. (1) Schematic of the standardized measuring area. (2) The custom 3D-printed holder securing the 2 × 3 OPM array. (3) The angle-adjustable design allows the sensors to be oriented perpendicular to the local surface contour of the muscle. **(C)** Experimental setup. Subjects were positioned supine on a custom non-magnetic platform. The left leg was secured with non-magnetic straps to prevent compensatory displacement during isometric ankle dorsiflexion. The inset illustrates the OPM sensor capturing the corresponding magnetic field generated by the neuromuscular activity along the z-axis. **(D)** Visual feedback system. A real-time trajectory tracking interface. Subjects dynamically adjusted their applied force (red line) to match the predefined target force trajectory (black line). Precise overlap indicates accurate execution of the ramp task. **(E)** Experimental paradigm and representative raw data. Top: The chronological protocol timeline. Following baseline noise recording and maximal voluntary contraction (MVC) determination, subjects performed five randomized submaximal ramp contractions (10%, 20%, 40%, 60%, and 80% maximal voluntary contraction, MVC). To assess repeatability, 20% and 60% MVC trials were subsequently repeated. All ramps featured a fixed rate of force development/relaxation (10% MVC/s) and a 10-s steady plateau. Rest intervals (1–5 min) were scaled to the preceding contraction intensity to prevent fatigue. Bottom: A continuous representative trace of the filtered OPM-MMG signal amplitude (pT) reflecting the sequential execution of the entire experimental paradigm.

To acquire TA MMG signals, a 2 × 3 array comprising 6 OPM sensors (FieldLine V2; FieldLine Inc., Boulder, CO, USA) was used. The sensors were housed in a custom polylactic acid holder, fabricated using a 3D printer (UltiMaker S7, UltiMaker, Netherlands). Within the holder, the inter-sensor spacing was set to 13.25 mm in the transverse direction and 22.00 mm in the longitudinal direction. Each mounting slot was dimensioned 1 mm larger than the OPM sensor casing to ensure a snug, movement-free fit (Figure 1B-1). For signal acquisition, the assembled OPM array was placed directly over the previously defined middle segment of the TA muscle. Specifically, the central longitudinal axis of the OPM array was aligned parallel to the line connecting the midpoints of the proximal and distal transverse tangents (Figure 1B-2). Furthermore, given the anatomical curvature of the anterior lower leg, the custom holder was designed with an adjustable central axis between the two columns of sensor slots. This allowed the two columns to rotate and conform to the curved surface of the leg, ensuring that both rows of OPM sensors remained as perpendicular as possible to the local skin surface to optimize magnetic signal detection (Figure 1B-3).

**Figure 2.**
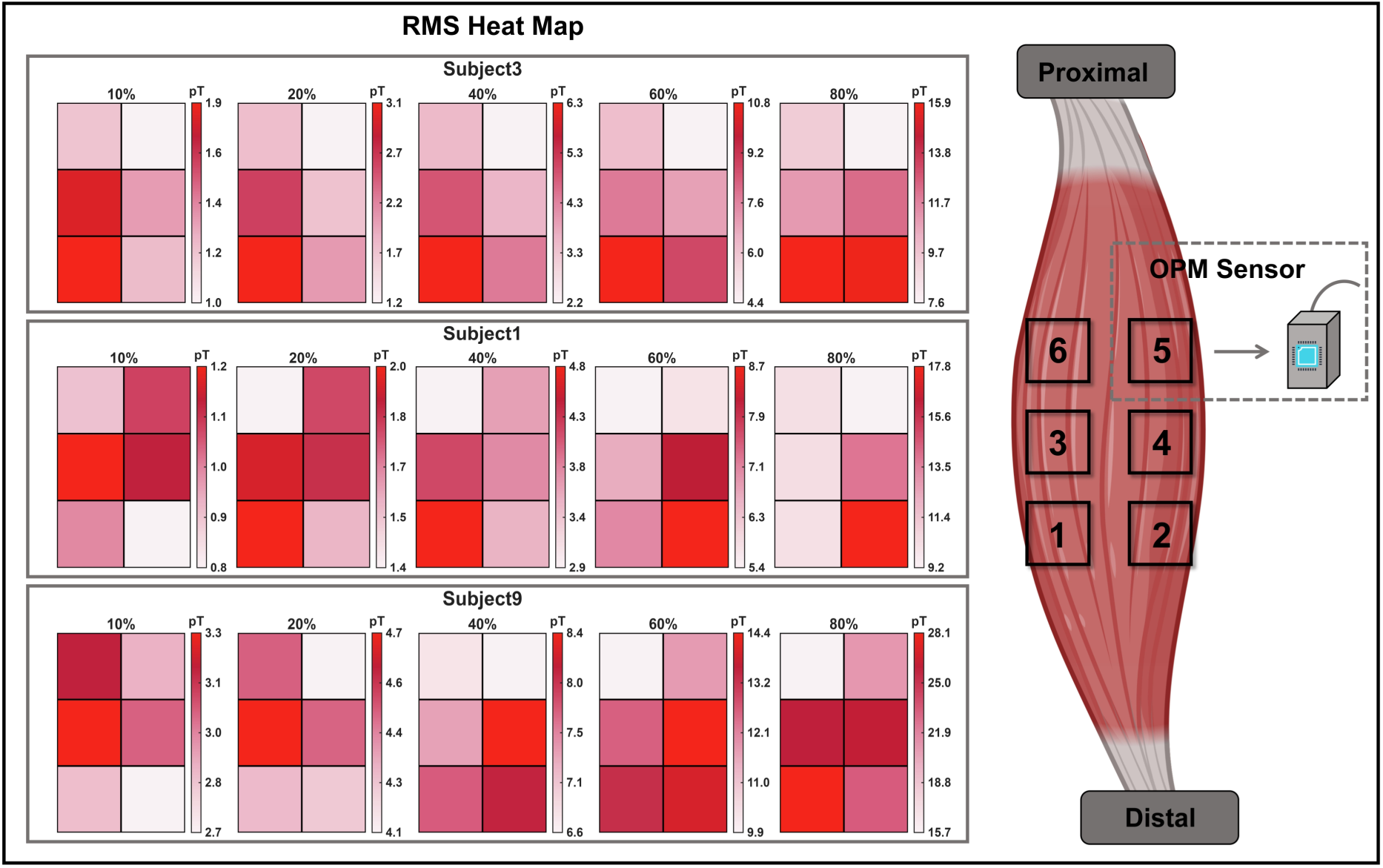
Spatial distribution of optically pumped magnetometer magnetomyography (OPM-MMG) root-mean-square (RMS) across varying isometric contraction intensities. (Left) RMS (i.e., the signal amplitude) heatmaps for three representative subjects (Subjects 3, 1, and 9) during the 10%, 20%, 40%, 60%, and 80% maximal voluntary contraction (MVC) tasks. Each 2 x 3 grid corresponds to the spatial arrangement of the optically pumped magnetometer (OPM) array. Within each subject and force level, the color scale was normalized to the local minimum (white) and maximum (dark red) RMS, so that the scaling remained relative and the visual pattern was preserved. (Right) Illustration of the 2 x 3 OPM sensor array positioned over the target muscle, providing the anatomical mapping (Sensors 1–6) for the corresponding heatmap grids. As depicted, Subject 3 exhibits a highly consistent spatial activation pattern across all intensities, whereas Subjects 1 and 9 show task-dependent shifts in the maximum RMS location.

**Figure 3:**
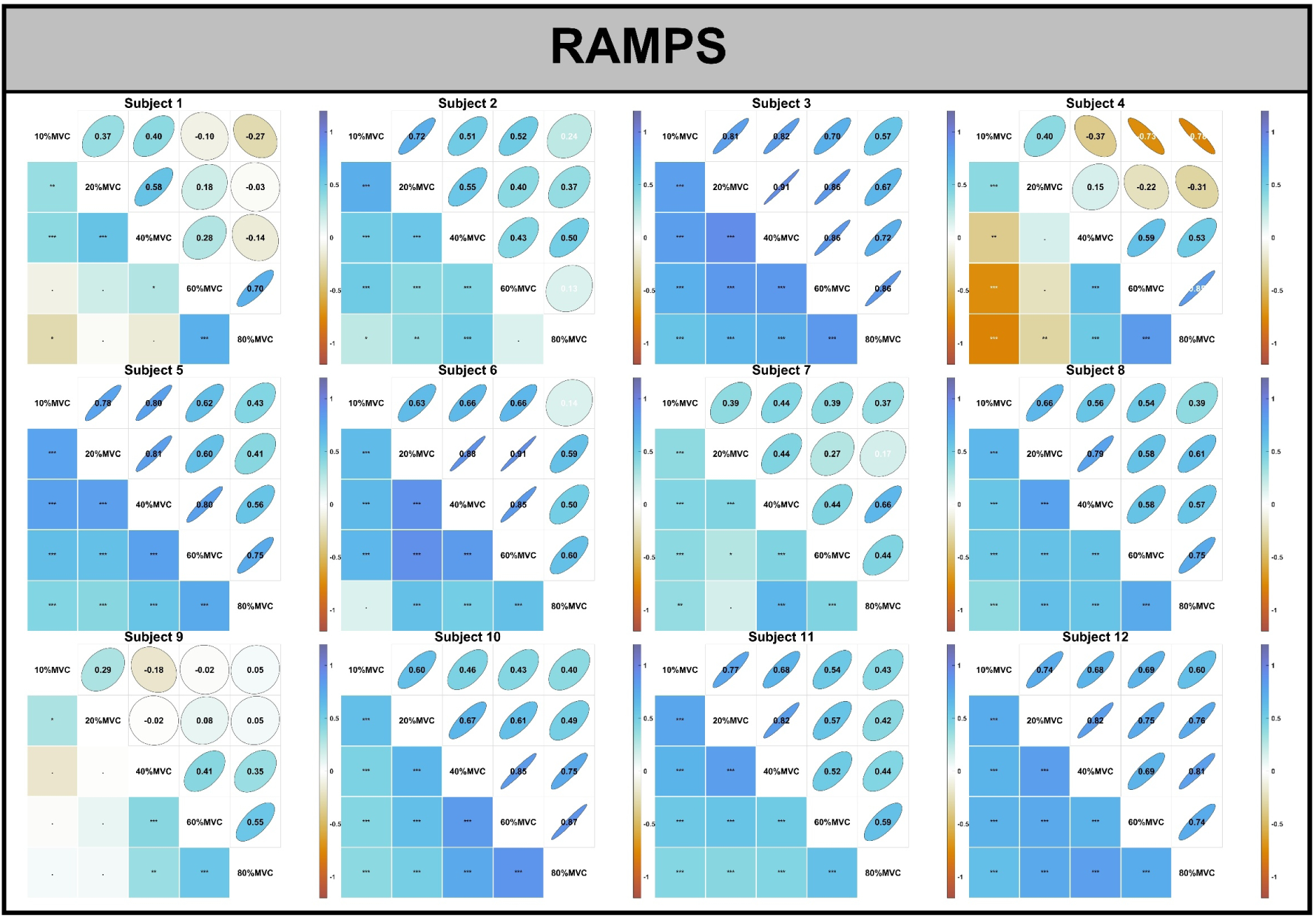
For each subject, Spearman’s rank correlation coefficients (ρ) were calculated among the 10%, 20%, 40%, 60%, and 80% maximal voluntary contraction (MVC) tasks using the dynamic root-mean-square (RMS) from the six OPM sensors across the 12 time windows. The upper triangular part of each matrix displays the correlation coefficients, whereas the lower triangular part indicates the corresponding significance levels. Positive ρ values indicate preservation of the relative spatial ordering of RMS across force levels, whereas negative values indicate an inverse ranking relationship. The diagonal shows the corresponding MVC tasks. (Significance is indicated as follows: **·** for p > 0.05, * for p < 0.05, ** for p < 0.01, and *** for p < 0.001).

During sensor placement, the six OPM sensors were sequentially inserted into the 2 × 3 slots of the 3D-printed holder to form the OPM array. The custom holder featured an angle-adjustable design between the two columns, which allowed the sensors to be oriented as perpendicular as possible to the local surface of the targeted TA region (Figure 1B-3). As dictated by the right-hand rule, longitudinal intracellular action currents propagating along the muscle fibers generate circulating magnetic fields (Figure 1C). By orienting the sensors’ sensitive z-axes approximately perpendicular to the local muscle surface, the array was positioned to record the magnetic field component along these axes, thereby improving sensitivity and quality of the OPM-MMG signals. Furthermore, according to the Biot-Savart law [25,26] and previous studies [8,9], the sensor-to-source distance significantly affects the recorded skeletal MMG signals. Therefore, we adjusted the distance between the OPM outer shell and the skin to be as minimal as possible while strictly avoiding direct physical contact (ranging from 6 to 9 mm during relaxation across all subjects). Specifically, this distance was minimized so that even during MVC — when muscle bulging and deformation are at their peak — the skin would not touch the OPM sensors (ranging from 1 to 3 mm during MVC across all subjects).

### 2.3 Experimental Design and Data Acquisition

After the ultrasound assessment, all subjects were comfortably positioned supine on a custom non-magnetic wooden platform (Figure 1C) inside a magnetically shielded room (Ak3b, VAC Vacuumschmelze, Hanau, Germany) at the MEG Center of the University of Tübingen, Germany, in June 2024. The left leg was aligned directly with the longitudinal axis of the force sensor. To prevent compensatory leg displacement during forceful contractions, the left leg was secured to the platform using non-magnetic straps at the ankle and thigh. The straps were tightened sufficiently to provide stable fixation without inducing passive muscle tension or involuntary contractions. A non-magnetic cord was secured around the foot, positioned just below the prominence of the first metatarsophalangeal joint. To ensure accurate force transmission, the distance between the subject’s foot and the force sensor was adjusted. Specifically, subjects moved longitudinally along the platform until the connecting cord showed neither slack nor active tension, with the ankle remaining in a neutral anatomical position. The experimental protocol commenced with a 1-minute baseline recording to record the resting-state noise levels. Subsequently, subjects performed three MVCs via isometric ankle dorsiflexion to maximally activate the TA muscle. Each MVC trial lasted 5 seconds, and a 30-second rest interval was provided between trials to prevent muscular fatigue.

Subsequently, isometric ramp contraction tasks at different force levels were performed based on each subject’s MVC. Five target levels were included: 10%, 20%, 40%, 60%, and 80% MVC. To prevent anticipation effects, subjects were blinded to the task sequence, which was completely randomized for each subject using a custom MATLAB (The MathWorks, Natick, MA, USA) script. Each ramp task consisted of three distinct phases: an ascending phase, a steady plateau phase, and a descending phase. To ensure a consistent rate of force development and relaxation across different target levels, the ramp rate was fixed at 10% MVC/s. In this way, we aimed to ensure that the observed variations in spatial distribution primarily reflected differences induced by the target force levels themselves, rather than being confounded by fluctuations in the rate of force development. Consequently, the duration of the ascending phase to reach the target force was 1 s for 10% MVC, 2 s for 20% MVC, 4 s for 40% MVC, 6 s for 60% MVC, and 8 s for 80% MVC. The descending phase mirrored the ascending phase in duration, and the steady plateau phase was maintained for 10 s across all target levels.

To prevent muscle fatigue from confounding the neuromuscular signals, customized rest intervals were implemented after each ramp task based on the preceding contraction intensity. Specifically, subjects rested for 1 min after the 10% MVC task, 2 min after the 20% MVC task, 3 min after the 40% MVC task, 4 min after the 60% MVC task, and 5 min after the 80% MVC task. After completing the initial block of five randomized ramp tasks, subjects were instructed to maintain their initial supine posture without any physical displacement and to rest quietly for 5 min. Subsequently, to assess the repeatability of the measurements, subjects performed a second block comprising the 20% and 60% MVC ramp tasks, with a strict 5-min rest interval between the two trials. To minimize confounding factors at extreme intensities, we restricted the repeatability analysis to 20% and 60% MVC. Excessively low levels (e.g., 10% MVC) elicit highly flexible force-generation strategies where the TA may not consistently dominate, whereas near-maximal levels (e.g., 80% MVC) introduce fatigue and synergistic co-activation of neighboring muscles that may complicate spatial pattern interpretation [24]. Therefore, 20% and 60% MVC were selected as more stable low-to-moderate and moderate-to-high load conditions, respectively, to evaluate within-subject spatial repeatability reliably. Finally, the experimental session concluded with a post-experiment 1-min baseline noise recording. The OPM signal outputs were continuously recorded directly by the FieldLine data acquisition system at a sampling frequency of 1000 Hz.

### 2.4 Force measurement device & visual feedback system

The force exerted during the experiment was measured using a custom-built non-magnetic device made of aluminum, equipped with strain gauge resistors (HBM, 1-LY41-6/350A, Resistance 350 Ω ± 0.3%, Gauge factor 2.07% ± 1%, transverse sensitivity −0.1%). To facilitate precise force application and real-time adjustments during the submaximal contractions, a visual feedback system was integrated with the force measurement setup. A monitor positioned in the subject’s field of view displayed a real-time graphical interface (Figure 1d). The interface displayed the predefined target force trajectory, including an ascending ramp, a steady plateau, and a descending ramp, represented by a solid black line. Simultaneously, the subject’s actual applied force was superimposed on the screen as a continuous red line. Subjects were instructed to dynamically adjust their contraction force to ensure the red line continuously tracked and overlapped the black target line as closely as possible. As illustrated in the feedback system, if the red line deviated above or below the black line, it indicated that the applied force was greater or less than the target, respectively. Precise overlap of the two lines demonstrated accurate force application.

### 2.5 Processing

All analyses were performed in MATLAB R2025b, the FieldTrip toolbox [27], and custom scripts. Raw OPM signal datasets were initially band-pass filtered between 10 Hz and 350 Hz using zero-phase, 4th-order IIR Butterworth filters. Powerline contamination and its harmonics were attenuated using a 4th-order IIR Butterworth band-stop filter (frequencies: 49–51, 99–101, 149–151, 199–201 Hz). At this point, filtered signals were visually inspected, and artifact-contaminated segments were manually annotated using interactive marking and later removed. Following artifact rejection, to quantify the signal amplitude, the root-mean-square (RMS) was calculated using 0.5-second non-overlapping windows, while the underlying datasets maintained their native sampling rate of 1000 Hz. For all subsequent analyses across the various MVC levels and repeated trials, data extraction was restricted to the 10-second steady-state plateau phase of each ramp task. Using custom MATLAB code, the initial and final 2 seconds of each plateau were trimmed to ensure the maximum stability of the RMS estimates. Consequently, the central 6-second segment — comprising 12 consecutive 0.5-second windows — was retained (i.e., from time window 1 to time window 12). This trimming procedure was implemented to isolate the most steady-state phase of the contraction, ensuring the analyzed data accurately represented the target MVC level by excluding any transitional force adjustments at the beginning and end of the task.

### 2.6 Statistical Analysis

Statistical analyses were performed to assess both the relationship of spatial MMG distributions across contraction intensities and the within-subject repeatability of these distributions during repeated measurements. All statistical analyses were conducted using MATLAB R2025b.

To determine whether the MMG activity pattern was preserved across force levels, pairwise comparisons were performed among the 10%, 20%, 40%, 60%, and 80% MVC tasks. For each subject and each MVC task, RMS values were extracted from 12 consecutive 0.5-s time windows at each of the six OPM recording positions, resulting in 72 RMS observations per MVC task. Each observation was defined by a specific OPM position and time-window index, and this order was kept identical across all MVC tasks. For example, OPM1-time window1 at 10% MVC was paired with OPM1-time window1 at 20% MVC, and OPM6-time window12 at 10% MVC was paired with OPM6-time window12 at 20% MVC. Within each MVC task, the 72 RMS observations were converted into ranks from 1 to 72. Spearman’s rank correlation coefficient (ρ) was then calculated between the corresponding rank sets for each force-level. Thus, this analysis assessed whether RMS observations with relatively high or low ranks at one force level retained similar relative ranks at another force level, rather than testing exact numerical agreement of RMS amplitudes. For all correlation analyses, p < 0.05 was considered statistically significant.

To assess within-subject spatial repeatability, the repeated 20% and 60% MVC ramp tasks were compared with their corresponding initial trials. For each subject, Spearman’s rank correlation coefficient was calculated to evaluate the preservation of the relative spatial ordering between the original and repeated measurements. In addition, intraclass correlation coefficients (ICC) were computed to assess the absolute numerical agreement between the two measurements. A two-way mixed-effects model for single measurements with absolute agreement (i.e., ICC (3,1)) was used [28]. For descriptive interpretation, ICC values below 0.50 were considered to indicate poor repeatability, values between 0.50 and 0.75 moderate repeatability, values between 0.75 and 0.90 good repeatability, and values above 0.90 excellent repeatability. Although ICC values are theoretically expected to range from 0 to 1, negative values may occasionally occur, indicating extremely poor agreement. Together, these complementary analyses allowed repeatability to be comprehensively evaluated from both rank-preservation and absolute-agreement perspectives.

ICC values were compared between the 20% and 60% MVC tasks across subjects. Normality of the paired ICC differences was assessed using the Shapiro-Wilk test. If the paired differences deviated significantly from normality, the nonparametric Wilcoxon signed-rank test was used. Statistical significance was defined as p < 0.05.

## 3. Results

The experiment was performed on 12 healthy subjects. All subjects and data were included in the final analysis. Of note, the subcutaneous fat thickness measured by ultrasound ranged from 3.1 to 5.2 mm across all subjects and was not statistically associated with the following results.

### 3.1 Spatial distribution of RMS across the OPM array at different contraction intensities

The consistency of the experimental setup was verified by post-session checks, during which the distance between the outer shell of the sensors and the skin remained 6–9 mm during relaxation and 1–3 mm during MVC across all subjects.

As expected, the mean RMS across all sensors generally increased with the targeted force level. Across all subjects and MVC tasks, the observed mean RMS ranged from 0.79 to 28.11 pT; the corresponding mean RMS are provided in Supplementary Table 1. To assess the spatial characteristics of the recorded MMG signals, heatmaps were generated to visualize the spatial distribution of mean RMS across the six OPM sensors at 10%, 20%, 40%, 60%, and 80% MVC (Figure 2). For each subject and MVC task, the color axis was scaled separately to the minimum and maximum mean RMS values across the six sensors in that specific panel. Thus, the color bars differ across MVC tasks because the local RMS minimum and maximum for each subject and force level defined the displayed color range. This panel-specific min–max color scaling was used to emphasize which OPM position showed the highest and lowest mean RMS within each MVC task, with red indicating the maximum and white the minimum. In this way, the dominant OPM location within the array’s spatial layout can be identified more intuitively through visual inspection. Consequently, the heatmaps are intended to illustrate the relative spatial ranking of mean RMS across the array within each force level, rather than to support direct color-based comparisons of absolute magnitude between different force levels.

Qualitative assessment of these RMS heatmaps revealed varying degrees of intra-subject spatial stability across the escalating contraction intensities. Figure 2 presents the spatial distribution patterns of three representative subjects (Subjects 3, 1, and 9) who demonstrated typical variations in spatial behavior. The remaining subjects’ RMS heatmaps are shown in Supplementary Figure 1. Although the specific OPM position recording the highest RMS varied among these individuals, each subject consistently maintained a distinct peak activation zone as force increased. For example, Subject 3 (Figure 2, top panel) demonstrated this consistent spatial pattern, where the maximum RMS localized predominantly over the distal-medial portion of the measurement zone (Sensors 1 and 3) across all force levels from 10% to 80% MVC.

Conversely, other subjects exhibited spatial shifts in the region of maximum RMS as force requirements increased. For instance, Subject 1 (Figure 2, middle panel) displayed peak activation primarily over the middle-left region (Sensor 3) at the lowest intensity (10% MVC). However, as the target force escalated to 60% and 80% MVC, the region of maximum RMS shifted toward the opposite aspect of the array, peaking at Sensors 2 and 4. Similarly, Subject 9 (Figure 2, bottom panel) showed spatial variation, with the peak amplitude transitioning between adjacent sensors within the lower and middle segments of the array as the target MVC changed.

### 3.2 Quantitative Assessment of Spatial Consistency

To quantitatively evaluate the spatial patterns observed in the RMS heatmaps, Spearman’s rank correlation analysis was performed. For each subject, correlation coefficients (ρ) were calculated for all pairwise comparisons among the 10%, 20%, 40%, 60%, and 80% MVC tasks (Figure 3). Overall, 9 of the 12 subjects exhibited varying degrees of positive spatial correlation across force levels. Across all subjects, positive correlations were more frequently observed between adjacent force levels, such as 10% versus 20% MVC or 40% versus 60% MVC, with peak ρ values of 0.91. In contrast, correlation coefficients generally decreased when more widely separated force levels were compared. For example, comparisons between 10% and 80% MVC yielded ρ values ranging from −0.78 to 0.60 across subjects.

At the individual level, the 9 subjects who showed stable activation patterns in the visually qualitative heatmap assessment generally demonstrated different positive correlations across most force-level comparisons (ρ = 0.13–0.91). Subject 3 is a representative example, showing predominantly positive correlations across the matrix (ρ = 0.57–0.91, p < 0.001). In this subject, strong positive correlations were observed both between adjacent force levels, such as 10% versus 20% MVC (ρ = 0.81, p < 0.001), and across broader force intervals, such as 20% versus 40% MVC (ρ = 0.91, p < 0.001). By contrast, subjects with visible spatial shifts in the heatmaps, including Subjects 1, 4, and 9, showed weaker, non-significant, or in some cases negative correlations, particularly when low-force conditions were compared with higher-force levels. For example, Subject 1 and Subject 9 showed correlations close to zero for the comparison between 10% and 80% MVC (ρ = −0.27 and ρ = 0.05, respectively; p > 0.05). In Subject 4, a strong negative correlation was observed across 10%-80% MVC (ρ = −0.78).

### 3.3 Within-subject repeatability of spatial MMG patterns

To evaluate within-subject repeatability at the two force levels (20% and 60% MVC), Spearman rank correlations (ρ) were calculated across the initial and repeated measurements for each of the 12 subjects (Figure 4). Specifically, for the correlations between the initial and repeated 20% MVC trials, with the sole exception of Subject 9, which showed a negative correlation (ρ = −0.45), the remaining 11 subjects exhibited positive correlations (ranging from ρ = 0.40 to 0.94). Similarly, for the comparisons between the initial and repeated 60% MVC trials, all 12 subjects consistently demonstrated positive correlations, ranging from ρ = 0.26 to 0.93. Furthermore, when comparing same-force versus across-force conditions, 7 of the 12 subjects yielded within-force correlations that were numerically higher than both mismatched cross-force comparisons (i.e., initial 20% vs. repeated 60% MVC (ρ = 0.23 to 0.87), and initial 60% vs. repeated 20% MVC (ρ = −0.42 to 0.84). The remaining five subjects exhibited more mixed patterns.

**Figure 4:**
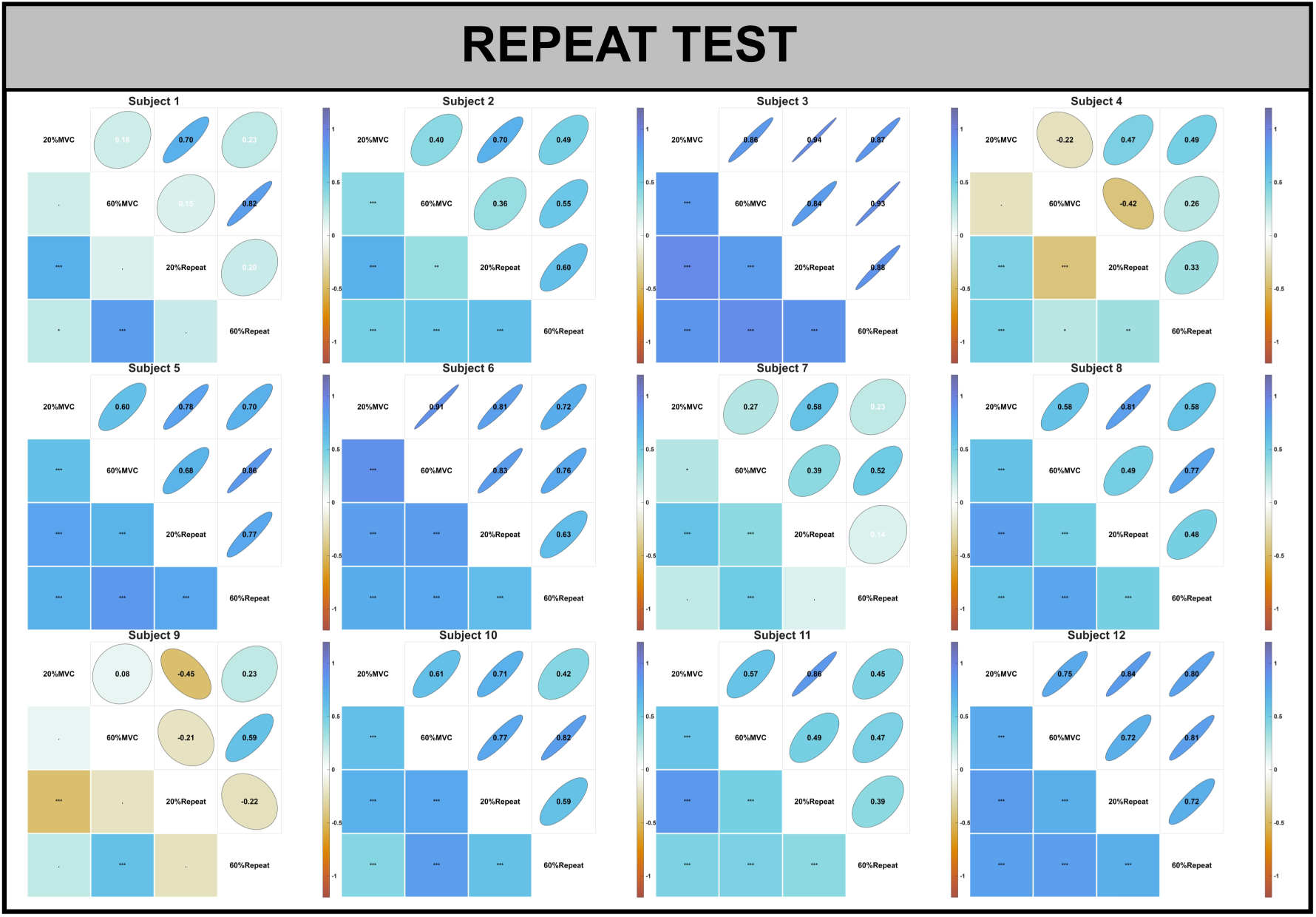
Within-subject repeatability of spatial Magnetomyography (MMG) patterns across initial and repeated force conditions. Spearman’s rank correlation matrices are presented for each of the 12 subjects. The comparisons encompass four experimental trials: the initial 20% maximal voluntary contraction (MVC), the initial 60% MVC, the repeated 20% MVC, and the repeated 60% MVC. The numerical values in the lower triangular cells represent the corresponding Spearman correlation coefficients (ρ). The magnitude and direction of these correlations are visually encoded by both the color gradient and the shape of the ellipses in the upper triangle. The color scale ranges from ρ = −1 (yellow/brown) to ρ = 1 (dark blue), with blue shades indicating positive correlations and yellow/brown shades indicating negative correlations. Narrower ellipses oriented diagonally (bottom-left to top-right) reflect stronger positive correlations, whereas more circular shapes represent weaker relationships.

Representative cases effectively illustrate this inter-subject variability. For instance, Subject 3 demonstrated consistently high positive correlations across all pairwise comparisons, displaying particularly robust repeatability for both the 20% (ρ = 0.94) and 60% MVC (ρ = 0.93) trials. Subject 1 exhibited a more selective pattern: correlations between the initial and repeated measurements at matched force levels were strong (20% MVC: ρ = 0.70; 60% MVC: ρ = 0.82), whereas the mismatched cross-force comparisons were substantially weaker (initial 20% vs. repeated 60% MVC: ρ= 0.23; initial 60% vs. repeated 20% MVC: ρ = 0.15). Conversely, Subject 9 presented the weakest repeatability, showing a negative correlation between the initial and repeated 20% MVC trials (ρ = −0.45), though the 60% MVC repeat comparison remained moderately positive (ρ = 0.59). Cross-force comparisons for this subject were similarly poor (initial 20% vs. repeated 60% MVC: ρ = 0.23; initial 60% vs. repeated 20% MVC: ρ = −0.21).

### 3.4 Absolute agreement of spatial MMG patterns

Having established the relative consistency of the spatial MMG patterns using Spearman rank correlations (Figure 4), the ICC (3,1) was subsequently employed to rigorously quantify their agreement between the initial and repeated measurements across all 12 subjects (Figure 5). Specifically, for the comparison between the initial and repeated 20% MVC trials, ICC values ranged from −0.2797 to 0.9588. Except for Subject 9 (ICC = −0.2797), the remaining 11 subjects exhibited positive agreement ranging from 0.3691 to 0.9588. Similarly, for the comparison between the initial and repeated 60% MVC trials, all 12 subjects demonstrated positive ICC values, with a narrower range from 0.2211 to 0.9252.

**Figure 5:**
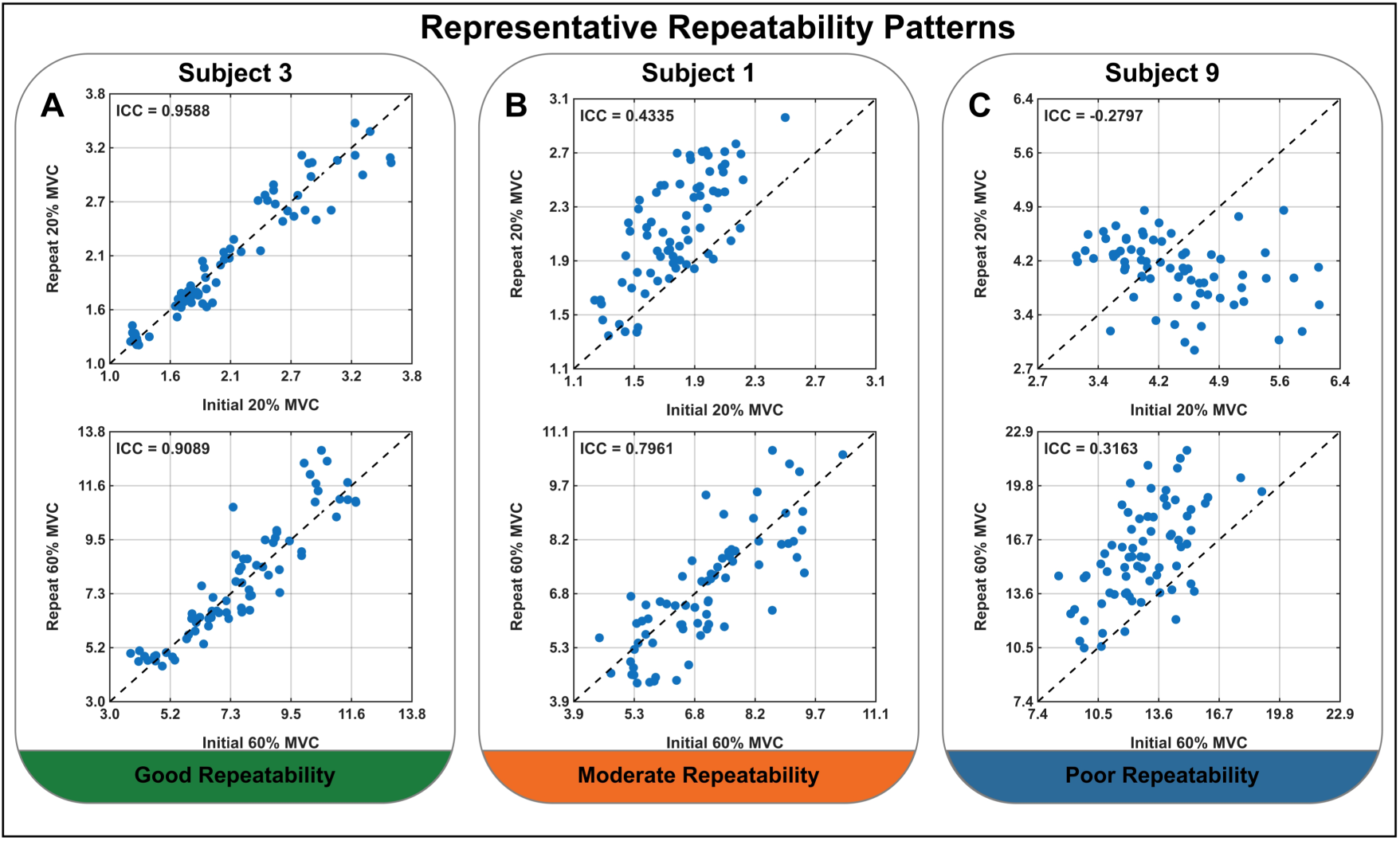
Root-mean-square (RMS) comparison of spatial Magnetomyography (MMG) patterns between initial and repeated measurements. The Intraclass Correlation Coefficient [ICC (3,1)] was calculated to strictly quantify within-subject repeatability at 20% and 60% MVC. Representative examples illustrate three distinct reliability profiles observed across the cohort: **(A)** Subject 3 demonstrates consistently high absolute agreement across both tested force levels. **(B)** Subject 1 exhibits a force-dependent pattern, with lower agreement at 20% MVC that notably improves at 60% MVC. **(C)** Subject 9 displays the weakest overall repeatability, recording a negative agreement at 20% MVC and maintaining a low level of agreement at 60% MVC. Detailed ICC results for the remaining subjects are provided in Supplementary Figure 2.

Representative examples further illustrate the specific patterns of agreement observed across individuals and force levels. Subject 3 (Figure 5A) demonstrated consistently high repeatability in both conditions, recording ICC values of 0.9588 for the repeated 20% MVC trials and 0.9089 for the repeated 60% MVC trials. Subject 1 (Figure 5B) exhibited a more moderate, force-dependent pattern, showing lower agreement between the initial and repeated 20% MVC trials (ICC = 0.4335), which notably increased for the repeated 60% MVC trials (ICC = 0.7961). Conversely, Subject 9 (Figure 5C) displayed the weakest overall repeatability, recording a negative agreement for the 20% MVC repeat trials (ICC = −0.2797) and maintaining a low level of agreement for the 60% MVC repeat trials (ICC = 0.3163). Detailed ICC results for the remaining subjects are provided in Supplementary Figure 2.

Across subjects, ICC values tended to be higher at 60% MVC than at 20% MVC. The median ICC was 0.6300 at 20% MVC and 0.8122 at 60% MVC, with corresponding mean values of 0.6122 and 0.6528, respectively. Nevertheless, the paired Wilcoxon signed-rank test showed no significant difference between the two force levels (p = 0.79 > 0.05).

## 4. Discussion

To our knowledge, this is the first study to investigate the characteristics and reliability of spatiotemporal MMG patterns in the TA across varying isometric force levels. The main results and conclusions are as follows:

● Consistent with our hypotheses, MMG spatial patterns were generally preserved across different force levels in most subjects (75%, i.e., 9 out of 12; Figure 3), although 25% of the subjects exhibited pronounced force-dependent spatial reorganization, which might be attributed mainly to physiological variability (see below for potential explanations).
● MMG spatial activity patterns are reliable when repeated, as shown by our repeated-task analyses at MVC levels of 20% (11 out of 12) and 60% (all subjects; see also Figures 4 and 5).

Together, these findings highlight the capability of OPM arrays in reliably characterizing individualized, force-dependent MMG activity, but also identify methodological challenges for future OPM array studies for MMG — in short, to optimize the reliability of OPM further, array-based MMG recordings, future studies should consider broader sensor coverage of the target muscle or muscle groups, real-time monitoring of limb movement, more precise control of sensor-to-source geometry, and complementary HDsEMG recordings for motor-unit-level interpretation of the observed MMG spatial patterns (see limitation section).

### 1) The RMS Heatmaps of MMG Activity

The RMS heatmaps visually showed spatial MMG activity variation across recording positions and force level (Figure 2). Specifically, half of the subjects (6 of 12) consistently showed the highest RMS at the same recording position across all target force levels (e.g., Subject 3), whereas the other subjects showed shifts in the recording position with the highest RMS (e.g., Subject 9). Although the position of the one-size-fits-all OPM array was standardized and the subjects’ lower legs were fixed, several factors may still have influenced the recorded RMS heatmaps of MMG activity: (1) the OPM array coverage may have included signal contributions from adjacent muscles; (2) subtle subject movements may have occurred during the measurement; (3) individual force-generation strategies may have differed across MVC tasks [29]; and (4) inter-subject differences in TA anatomy may have led to different contraction-related changes in sensor-to-source orientation and distance. Specifically, these factors can be elaborated as follows: (1) Due to inter-subject variations in TA muscle size, the coverage of the fixed-size OPM array may encompass adjacent muscles, such as the extensor digitorum longus [30]. Particularly during high-intensity tasks (e.g., 80% MVC), these synergistic muscles may co-activate to achieve the target force, leading to signals that may not exclusively reflect TA activity. (2) Although the subjects’ lower legs were securely fixed, slight, unintended movements could still occur during isometric contractions. This may cause the predefined TA target area (i.e., the signal source) to shift relative to the OPM array, leading to a spatial deviation of the signal source. (3) Force generation strategies across different MVC tasks vary among individuals. For instance, in Subjects 7, 10, and 11, the location of the maximum RMS exhibited phase-specific characteristics: the highest RMS occurred at one consistent recording position during low-intensity tasks (10% and 20% MVC) and shifted to another stable location as the force increased (40%, 60%, and 80% MVC). In addition, during low-force tasks such as 10% MVC, only minimal effort was required to reach the target force, and the TA may not have been sufficiently activated as the dominant contributor to force production. As the MVC task increased, the force-generation strategy may have changed toward greater TA involvement, which could explain the shift of the highest-RMS region in the RMS heatmap. (4) From an anatomical perspective, the TA (i.e., the primary muscle responsible for ankle dorsiflexion) is a pennate muscle that undergoes individualized, non-uniform dynamic deformation (e.g., muscle bulging and fiber shortening) during isometric contractions. These structural changes dynamically alter the internal orientation between the active muscle fibers (i.e., the signal source) and the OPM sensors. In contrast to the more established application of MEG, where the signal source remains static, spatial variations are typically addressed using localization coils, enabling precise, real-time three-dimensional tracking [31]. OPM-MMG currently lacks comparable real-time signal source tracking technology [9]. Consequently, these unmonitored, subject-specific orientation shifts during contraction may contribute to the observed variations in the RMS heatmaps. In addition to these orientation variations, dynamic muscle deformation also alters the sensor-to-source distance. Instead of monitoring the sensor-to-source distance, we recorded the distance between the OPM sensor’s outer shell (i.e., the external casing) and the skin (maximum difference between pre– and post-session: 1 mm), which indirectly reflects changes in the underlying internal geometry. Since our previous research demonstrated that variations in this scale do not compromise OPM-MMG reliability [8], minor variations in the outer-shell-to-skin distance are unlikely to account for the observed MMG spatial differences.

### 2) Force-dependent preservation and reorganization of TA MMG spatial patterns

This MMG spatial pattern shift (Figure 3) could be interpreted through two mechanisms: motor unit recruitment and force-dependent changes in muscle geometry (e.g., muscle fiber shortening and muscle belly bulging), which alter the sensor-to-source relationship. First, between adjacent force levels (e.g., 10% and 20% MVC), the same motor units might remain active and exhibit similar firing rates, thereby preserving the spatial pattern of MMG activity, consistent with a previous HDsEMG study [23]. However, to verify this motor-unit-based interpretation, motor unit decomposition would be required, which is typically performed using HDsEMG recordings [32,33]. Specifically, the motor unit decomposition could identify which motor units are active at each force level and quantify whether their firing rates remain similar or change as force demand increases [34,35]. Since the present study did not include this reference modality (i.e., no simultaneous HDsEMG), changes in motor unit recruitment and firing rate should be regarded as plausible but speculative contributors to the observed MMG spatial pattern shifts. Compared with the adjacent force level, larger force spans (e.g., 10% to 80% MVC) might require recruiting additional motor units and further increasing firing rates. These changes could alter the relative contribution of signal sources within the TA and adjacent synergistic muscles (e.g., the extensor digitorum longus) to the recorded MMG signal, thereby shifting the spatial RMS ranking observed at lower force levels. Second, pronounced muscle fiber shortening and bulging at high force levels dynamically alter the sensor-to-source geometry, i.e., the orientation between the signal source and the OPM sensor, which may contribute to the observed shift in MMG spatial patterns.

Notably, 3 subjects (Subjects 1, 4, and 9) exhibited weak or negative correlations across distant force levels (e.g., 10% vs. 80% MVC in Subject 1, ρ = −0.27). To determine whether anatomical variations contributed to this discrepancy, we investigated subcutaneous tissue thickness; however, these differences could not be attributed to subcutaneous tissue thickness, as their values (0.36–0.49 cm) fell well within the cohort range (0.36–0.52 cm) [36]. Regarding the weak or negative correlations in MMG spatial patterns observed in such subjects across different MVC tasks, we speculate on two possible explanations: First, methodologically, this may result from deviations in the signal source caused by slight left leg movements during the experiment. Particularly during lower-target-force tasks requiring minimal effort, the TA might not be the dominant MMG signal source. Under these conditions, minor limb movements could easily cause geometric shifts [37–39], misaligning the recorded signals from the predefined TA area. For example, Subject 9 showed very weak or no spatial correlation even between similar low-intensity tasks (10% vs. 20% MVC, ρ = 0.29, p < 0.05; 20% vs. 40% MVC, ρ = −0.02, p > 0.05). Although the thigh was fixed to maintain signal-source stability, such minor movements are difficult to prevent without real-time monitoring. Notably, Subject 9’s outer-shell-to-skin distance remained unchanged across sessions (7 mm at rest, 3 mm during MVC). This indirectly confirms the OPM array’s physical stability relative to the measured TA area, suggesting that the signal-source shift was more likely related to relative limb movement than to displacement of the OPM array. Second, from a physiological perspective, we speculate that these differences in MMG spatial patterns may be related to changes in motor unit recruitment strategy and discharge behavior across force levels. For instance, Subjects 1 and 4 exhibited negative spatial correlations across different force levels (e.g., 10% vs. 60% or 80% MVC; ρ = −0.10, −0.27, and −0.73, −0.78, respectively), but strong positive correlations between adjacent high-force tasks (60% vs. 80% MVC; ρ = 0.70 and 0.85, respectively; p < 0.001). Accordingly, these weak or negative MMG spatial correlations are more likely due to a combination of signal-source variation and individualized neuromuscular control adjustments. Lacking real-time movement tracking and simultaneous HDsEMG for motor unit decomposition as a reference [40], both explanations should currently be interpreted cautiously as potential mechanisms.

Another notable finding is that in Subjects 7, 10, and 11, the recording position with the maximum RMS was not identical across all MVC tasks. However, these subjects still showed positive spatial correlations across force levels (Subject 7: ρ = 0.17– 0.66; Subject 10: ρ = 0.40–0.87; Subject 11: ρ = 0.42–0.82). This indicates that a shift in the highest recording position of the RMS heatmap (Supplementary Figure 1) does not necessarily imply a complete reorganization of the entire MMG spatial pattern. Spearman correlation reflects the overall preservation of the relative spatial distribution across recording positions, rather than the stability of a single maximum-RMS site. Therefore, in these subjects, the MMG spatial pattern may be better interpreted as partially preserved but force-dependently redistributed, with the dominant RMS region shifting as the target force increased, while the overall spatial organization remained partially preserved.

In summary, the results of the MVC tasks indicate that as the target force increases, the spatial MMG patterns of the TA neither change entirely randomly nor remain static. For the majority of subjects (9 out of 12), the spatial patterns across different force levels exhibit continuity, indicating that the fundamental spatial pattern of MMG signals is preserved. Simultaneously, as the force level increases, this pattern shifts progressively due to increased motor unit recruitment and contraction-induced changes in muscle morphology. For a minority of subjects, this shift is more pronounced and can even manifest as a significant reorganization of the spatial pattern. Overall, these findings demonstrate that the OPM array can reveal the diverse spatial MMG distribution characteristics exhibited by individuals in response to increasing force levels, thereby providing an approach for understanding the underlying differences in personalized neuromuscular control strategies.

### 3) Within-subject repeatability under repeated MVC tasks

The repeated measurements showed that MMG spatial patterns were preserved under the same MVC tasks. Specifically, for the comparison between the initial and repeated 20% MVC trials, 11 of 12 subjects showed positive Spearman correlations, ranging from ρ = 0.40 – 0.94, with only Subject 9 showing a negative correlation (ρ = −0.45). For the initial and repeated 60% MVC trials, all subjects showed positive correlations, ranging from ρ = 0.26 – 0.93. These results demonstrate that MMG spatial patterns were generally reproducible when the target force level was held constant, and the differences observed across different MVC tasks could be attributed to changes in target force. In some subjects (e.g., Subject 3), this repeatability was manifested not only in the preservation of the relative MMG spatial patterns but also in the high consistency (ICC = 0.9588 for 20% MVC and ICC = 0.9089 for 60% MVC) of localized RMS, suggesting that the motor unit recruitment patterns and muscle firing rates may have remained relatively stable [41]. In contrast, Subjects 1 and 9 showed weaker repeatability (ICC = 0.4335 and ICC = −0.2797, respectively) during the 20% MVC task but exhibited noticeable improvements during the 60% MVC task (ICC = 0.7961 and ICC = 0.3163, respectively). We speculate that the neuromuscular system may exhibit different motor unit recruitment flexibility during a low-MVC task. However, it should be noted that while a higher force level may result in more pronounced motor unit recruitment, it also induces greater muscle bulging and changes in the sensor-to-source relationship (e.g., distance and orientation), which can lead to local RMS variations. Moreover, this also helps explain the coexistence of high Spearman correlations with low ICC values observed in OPM-MMG measurements: the subjects reproduced the MMG spatial pattern, whereas RMS at corresponding recording positions showed less agreement between repeated measurements. Moreover, the statistical finding of no significant difference in ICC between the 20% and 60% MVC groups (p > 0.05) further confirms that this influence is highly subject-dependent.

Another notable observation from the repeated tests is that the cross-MVC-task spatial similarity (i.e., the spatial correlation between 20% and 60% MVC) showed a high degree of intra-subject consistency (e.g., for subject 3: the Spearman’s rank correlation between 20% MVC and 60% repeat is 0.87). Specifically, if a subject demonstrated cross-MVC-task spatial pattern similarity in the initial task, this characteristic was preserved (9 out of 12 subjects) in the repeated session (e.g., initial 20% MVC vs. repeated 60% MVC). Conversely, if MMG spatial pattern differences (p < 0.05) existed between the two MVC tasks initially, these differences were consistently reproduced in the repeated test (p < 0.05) rather than being attenuated (Figure 4). Therefore, lower cross-MVC-task similarity should not be misinterpreted as measurement error or poor data quality; rather, it reflects a distinctly different neuromuscular control strategy. A higher similarity suggests that the subject maintained a relatively stable MMG spatial pattern across force levels, possibly by accommodating the increased force demand mainly through broader increases in discharge rate and recruitment within an already active motor-unit pool. Conversely, lower similarity indicates that, when adapting to changes in the MVC task, the subject may rely more on spatial reorganization of local motor unit recruitment — that is, shifting the dominant activation zones to different anatomical regions of the muscle. Accordingly, the repeatability of these between-task differences provides support for the point that such MMG pattern differences represent subject-specific features of neuromuscular control rather than random variability.

Overall, relying solely on Spearman’s rank correlation may overestimate the spatial stability for certain individuals, as the preservation of relative spatial rankings does not equate to RMS agreement [42,43]. Conversely, relying exclusively on the ICC might underestimate pattern stability when the overall signal amplitude is entirely scaled while the spatial structure remains intact [28]. Therefore, the combined application of Spearman’s correlation and the ICC facilitates a more comprehensive interpretation of MMG spatial repeatability across repeated MVC tasks along two complementary dimensions: pattern preservation and RMS agreement. This approach not only characterizes the reproducibility of MMG spatial patterns under fixed tasks but also demonstrates certain methodological advantages: as a contactless measurement technique, it can provide spatially distributed information comparable to that obtained in previous HDsEMG studies [44,45] while avoiding the need for extensive skin preparation and electrode attachment, which may be advantageous for repeated experiments and future clinical applications.

## 5. Limitations and Perspective

Several limitations of the present study should be considered when interpreting the results. First, the 2 × 3 six-channel OPM array used in this study had limited coverage, recording only a localized region of the TA rather than the entire muscle. Therefore, the observed MMG spatial patterns should be interpreted as regional within the measured area rather than as a complete representation of the TA as a whole. Moreover, to better elucidate the underlying neuromuscular control strategies, future studies should include HDsEMG as a comparative tool to interpret these spatial results at the motor-unit level precisely. Second, the sample size was relatively small (N = 12). Although the current results preliminarily demonstrate that the OPM array can record regional spatial activation features of the TA, larger sample sizes are required to validate these findings further and comprehensively explore inter-individual differences. Future studies could use arrays comprising more OPM sensors to expand muscle coverage and characterize spatial MMG patterns more comprehensively. Furthermore, although sensor placement was standardized as much as possible, slight variations in sensor position, sensitive axis orientation, and outer shell-to-skin distance may still affect the MMG signals. While we strictly controlled for these factors during the experiment, they were not independently quantified; future research should conduct more detailed evaluations of these parameters. Therefore, future MMG studies could incorporate real-time monitoring of limb movement better to interpret MMG spatial pattern variations across different force levels. In addition, the repeatability analysis was only performed at two target force levels (20% and 60% MVC), and these findings cannot be directly extrapolated to other contraction intensities (e.g., 10%, 40%, or 80% MVC). Meanwhile, this study focused on the relatively superficial TA and employed a controlled isometric ramp contraction task. Therefore, further research is needed to determine whether the current conclusions can be generalized to deeper muscles, more complexly morphed muscles, or muscles that undergo significant displacement during dynamic tasks.

Despite these limitations, this study still provides a preliminary reference for the feasibility and reliability of OPM-MMG for assessing spatial patterns of local muscle activity and offers a basis for future study. First, regarding physiological mechanisms and clinical exploration, this study demonstrates that the OPM array not only effectively records neuromuscular activity but also reflects spatial pattern differences across recording sites, along with their dynamic characteristics as force levels change. Compared to a single OPM sensor during MMG study, the OPM array can characterize localized motor unit recruitment patterns and inter-individual neuromuscular control strategies in more detail. For diseases characterized by local neuromuscular dysfunction, this spatial and dimensional information can provide a valuable complementary perspective. Second, regarding translational applications, the current results suggest the potential utility of array-based measurements, and it is promising to extend this approach to more complex dynamic tasks and neuromuscular function assessments in a wider range of clinical scenarios [46].

## 6. Conclusion

This study demonstrates that a 2×3 array-based OPM-MMG is an applicable tool for mapping force-dependent MMG activity patterns and recording subject-specific spatial variability within localized muscles. By establishing this methodological basis, OPM-array-based MMG could be used in future neuromuscular research and may support the further translation of MMG toward clinical applications.

## 7. Declarations

### 7.1 Data availability statement

The original data can be made accessible upon reasonable request directed to the corresponding author.

### 7.2 Ethics statement

The study was conducted in accordance with the Declaration of Helsinki (World Medical Association, 2001) and approved by the ethics committee of the University of Tübingen. The subjects of this publication gave informed consent to publish their data.

### 7.3 Author contributions

HY and JM: study design, data collection and analysis, writing, and revision. BK: study design, data collection, and revision. BS: 3D printed models, data collection, and revision. All authors contributed to the article and approved the submitted version.

## 7.4 Acknowledgment

All authors would like to acknowledge the support of Jürgen Dax for providing the visual feedback system and force transducer used in this study.

## Abbreviations

MMG: Magnetomyography
OPM: Optically pumped magnetometer
OPM-MMG: Optically pumped magnetometer magnetomyography
TA: Tibialis anterior
MVC: Maximal voluntary contraction
SQUID: Superconducting quantum interference device
MEG: Magnetoencephalography
MCG: Magnetocardiography
sEMG: Surface electromyography
HDsEMG: High-density surface electromyography
RMS: Root-mean-square

---

**Supplementary Table 1.** Mean root-mean-square (RMS) values are reported for each subject and optically pumped magnetometer (OPM) recording position during the central 6 s of the plateau phase at 10%, 20%, 40%, 60%, and 80% MVC. Values are given in pT.

|  | Subject ID | OPM 1(pT) | OPM 2(pT) | OPM 3(pT) | OPM 4(pT) | OPM 5(pT) | OPM 6(pT) |
| --- | --- | --- | --- | --- | --- | --- | --- |
| 10% MVC (mean RMS) | Subject 1 | 0.9690 | 0.7925 | 1.1551 | 1.0913 | 1.0469 | 0.8879 |
|  | Subject 2 | 1.4969 | 1.5196 | 2.1811 | 1.7773 | 1.8677 | 2.2543 |
|  | Subject 3 | 1.8555 | 1.2588 | 1.7908 | 1.3601 | 1.0185 | 1.2271 |
|  | Subject 4 | 1.7395 | 1.5275 | 2.0172 | 1.2954 | 1.2454 | 2.1483 |
|  | Subject 5 | 1.3469 | 1.7193 | 1.3909 | 1.5531 | 1.3290 | 1.2177 |
|  | Subject 6 | 1.7095 | 2.0978 | 1.7622 | 2.0883 | 1.9896 | 1.6100 |
|  | Subject 7 | 1.3061 | 1.4141 | 1.6343 | 1.5328 | 1.5946 | 1.5000 |
|  | Subject 8 | 0.9039 | 1.0045 | 1.2161 | 1.3296 | 1.1559 | 1.4879 |
|  | Subject 9 | 2.8296 | 2.6581 | 3.2747 | 3.0287 | 2.8648 | 3.1647 |
|  | Subject 10 | 1.1690 | 1.2525 | 1.3937 | 1.3999 | 1.2219 | 1.0736 |
|  | Subject 11 | 0.7980 | 0.8098 | 1.1400 | 1.0684 | 1.0705 | 0.9667 |
|  | Subject 12 | 1.1680 | 1.1498 | 1.7430 | 1.5343 | 1.2004 | 1.2483 |
| 20% MVC (mean RMS) | Subject 1 | 1.9724 | 1.5812 | 1.9194 | 1.8411 | 1.7914 | 1.4035 |
|  | Subject 2 | 2.7252 | 2.5766 | 3.4345 | 2.9448 | 3.0947 | 3.4880 |
|  | Subject 3 | 3.1422 | 2.0464 | 2.5821 | 1.7386 | 1.2482 | 1.7694 |
|  | Subject 4 | 2.5694 | 2.5086 | 2.7401 | 2.5634 | 2.4010 | 3.1940 |
|  | Subject 5 | 1.8841 | 2.6506 | 1.8970 | 2.2390 | 1.7819 | 1.6177 |
|  | Subject 6 | 2.6395 | 5.1888 | 2.4693 | 3.8832 | 3.4279 | 2.7283 |
|  | Subject 7 | 2.2539 | 2.6221 | 2.9517 | 2.6369 | 2.9454 | 2.7656 |
|  | Subject 8 | 1.5730 | 2.0276 | 2.1688 | 2.1563 | 1.6761 | 2.6855 |
|  | Subject 9 | 4.3178 | 4.2794 | 4.7290 | 4.4941 | 4.1455 | 4.4984 |
|  | Subject 10 | 1.9621 | 2.3181 | 2.4041 | 2.5142 | 1.9645 | 2.1023 |
|  | Subject 11 | 0.9101 | 0.9511 | 1.3528 | 1.1776 | 1.2344 | 1.1616 |
|  | Subject 12 | 1.9297 | 2.0940 | 3.0856 | 2.5859 | 1.8627 | 2.0038 |
| 40% MVC (mean RMS) | Subject 1 | 4.7687 | 3.5109 | 4.1752 | 3.7922 | 3.6483 | 2.8989 |
|  | Subject 2 | 4.5180 | 5.6544 | 6.3402 | 6.3258 | 6.4811 | 6.7674 |
|  | Subject 3 | 6.3412 | 4.4097 | 4.8611 | 3.5287 | 2.2430 | 3.4603 |
|  | Subject 4 | 4.6068 | 5.5886 | 4.7202 | 5.7077 | 5.4455 | 5.3022 |
|  | Subject 5 | 4.6153 | 7.5385 | 3.6222 | 5.8266 | 3.7039 | 3.3215 |
|  | Subject 6 | 5.4302 | 11.5398 | 4.8948 | 8.2240 | 7.2836 | 5.2277 |
|  | Subject 7 | 4.2908 | 5.2404 | 5.2617 | 5.8850 | 6.1657 | 4.9897 |
|  | Subject 8 | 2.7855 | 3.7126 | 3.7350 | 3.6649 | 2.8736 | 5.3194 |
|  | Subject 9 | 7.7266 | 8.0730 | 7.3321 | 8.4141 | 6.6304 | 6.7830 |
|  | Subject 10 | 4.0090 | 5.6235 | 4.3253 | 5.1910 | 3.6643 | 3.8481 |
|  | Subject 11 | 1.3724 | 1.5227 | 1.9380 | 1.6407 | 1.9630 | 1.8928 |
|  | Subject 12 | 2.6135 | 3.3009 | 4.5086 | 3.5724 | 2.4866 | 2.5931 |
| 60% MVC (meanRMS) | Subject 1 | 7.0198 | 8.7272 | 6.6015 | 8.1839 | 5.7406 | 5.4343 |
|  | Subject 2 | 11.0385 | 12.2305 | 13.2044 | 12.0521 | 12.0606 | 15.1176 |
|  | Subject 3 | 10.8323 | 8.7857 | 7.7970 | 6.9379 | 4.3766 | 6.1802 |
|  | Subject 4 | 7.8082 | 10.7825 | 5.8157 | 12.7327 | 11.6858 | 8.8995 |
|  | Subject 5 | 7.5541 | 11.5746 | 5.8658 | 10.7444 | 7.0777 | 6.8365 |
|  | Subject 6 | 5.7253 | 7.0956 | 5.5882 | 6.6132 | 5.5018 | 5.5807 |
|  | Subject 7 | 6.3829 | 7.7574 | 7.4037 | 7.6543 | 7.8055 | 6.7947 |
|  | Subject 8 | 4.5089 | 4.2480 | 6.5022 | 4.4891 | 4.1638 | 8.9903 |
|  | Subject 9 | 13.3456 | 13.9391 | 12.6153 | 14.3536 | 11.7382 | 9.9094 |
|  | Subject 10 | 6.3889 | 9.6167 | 6.6813 | 7.5541 | 5.0103 | 5.3913 |
|  | Subject 11 | 2.7107 | 3.3010 | 3.4210 | 3.1242 | 3.7637 | 3.0248 |
|  | Subject 12 | 3.6039 | 4.7915 | 6.7079 | 5.3231 | 4.6491 | 5.2306 |
| 80% MVC (mean RMS) | Subject 1 | 10.1339 | 17.7588 | 10.2973 | 13.8333 | 9.2298 | 10.1857 |
|  | Subject 2 | 11.4175 | 14.1016 | 13.3077 | 15.1436 | 14.9368 | 16.8417 |
|  | Subject 3 | 15.8880 | 15.7785 | 11.0861 | 12.3555 | 7.6070 | 9.3765 |
|  | Subject 4 | 15.0390 | 19.1084 | 8.8754 | 24.2939 | 20.9490 | 14.5099 |
|  | Subject 5 | 8.0129 | 14.3846 | 5.9358 | 9.9831 | 7.4134 | 8.7335 |
|  | Subject 6 | 13.1233 | 27.7153 | 14.2171 | 17.8083 | 12.5499 | 18.4376 |
|  | Subject 7 | 10.3573 | 12.0832 | 11.4909 | 12.2955 | 12.5818 | 10.0138 |
|  | Subject 8 | 5.9094 | 5.1170 | 8.3117 | 5.1490 | 4.4681 | 12.7627 |
|  | Subject 9 | 28.1197 | 23.3700 | 26.0151 | 26.1673 | 21.0231 | 15.6699 |
|  | Subject 10 | 13.1784 | 15.6496 | 13.1622 | 14.1982 | 9.7480 | 9.6197 |
|  | Subject 11 | 5.3961 | 6.8668 | 6.6639 | 7.2881 | 8.9967 | 5.6055 |
|  | Subject 12 | 4.3885 | 7.6905 | 8.6235 | 8.1140 | 5.6263 | 5.7809 |

**Supplementary Figure 1.**
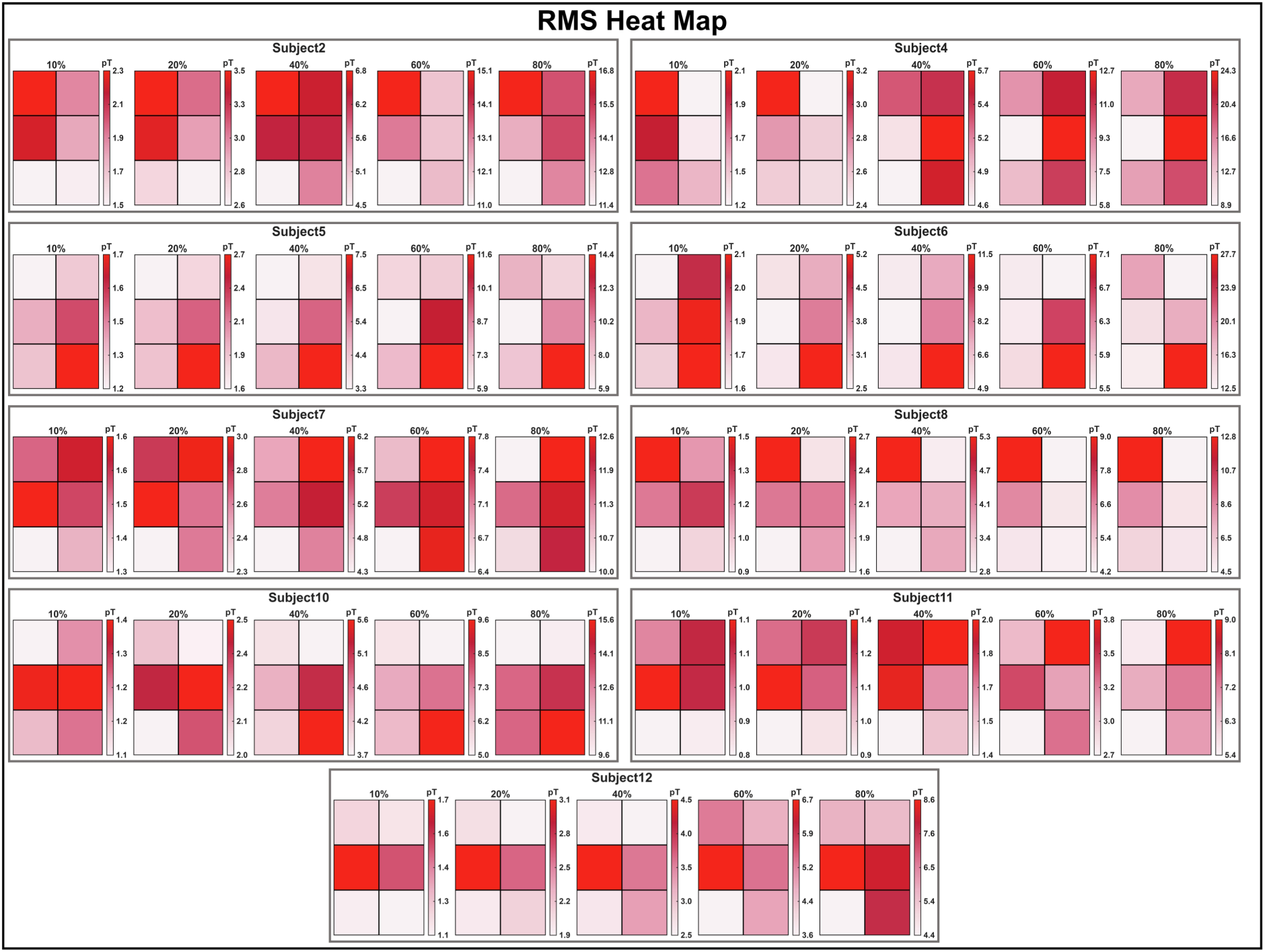
Root-mean-square (RMS) heatmaps of the remaining subjects are presented across the five maximal voluntary contraction (MVC) tasks (10%, 20%, 40%, 60%, and 80% MVC).

**Supplementary Figure 2.**
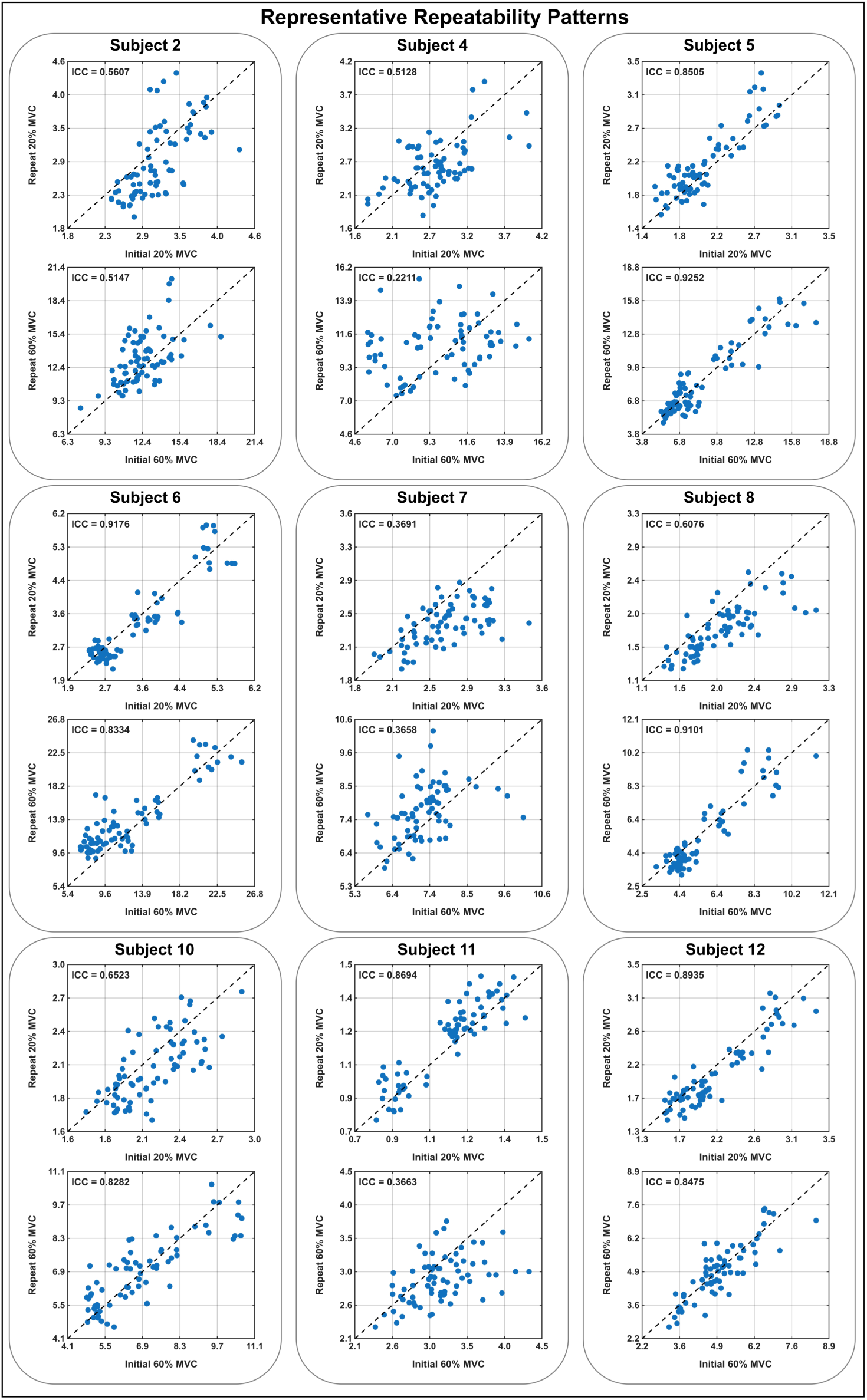
Root-mean-square (RMS) comparison of spatial Magnetomyography (MMG) patterns between initial and repeated measurements. The Intraclass Correlation Coefficient [ICC (3,1)] was calculated to strictly quantify within-subject repeatability at 20% and 60% maximal voluntary contraction (MVC) for the remaining subjects.

